# Grip force adjustments reflect prediction of dynamic consequences in varying gravitoinertial fields

**DOI:** 10.1101/190983

**Authors:** Olivier White, Jean-Louis Thonnard, Philippe Lefèvre, Joachim Hermsdörfer

## Abstract

One remarkable capacity when we grasp and manipulate tools relies on the ability to predict the grip force required to handle them in relation to their mechanical properties and the surrounding environment. However, rapid changes in the dynamical context may constitute a substantial challenge. Here, we test how participants can switch between different and never experienced dynamical environments induced by centrifugation of the body. Seven subjects lifted an object four times in a row successively in 1, 1.5, 2, 2.5, 2, 1.5 and 1g. We continuously measured grip force, load force and the gravitoinertial acceleration that was aligned with body axis (perceived gravity). Participants adopted stereotyped grasping movements immediately upon entry in a new environment and needed only one trial to adapt grip forces to a stable performance in each new gravity environment. While participants predictively applied larger grip forces when they expected increasing gravity steps, they did not decrease grip force proportionally when they expected decreasing gravity steps, indicating imperfect anticipation in that condition. The subjects’ performance could rather be explained by a combination of successful scaling of grip force according to gravity changes and a separate safety factor. The data suggest that in highly unfamiliar dynamic environments, grip force regulation is characterized by a combination of a successful anticipation of the experienced environmental condition, a safety factor reflecting strategic response to uncertainties about the environment and rapid feedback mechanisms to optimize performance under constant conditions.

## Introduction

Consider a worker whose job is to sort packages of varying size and weight into bins, bags, or slots. Each of these packages will have different inertial properties and will impose different loads on the arm. The dynamics of the upcoming movement is not fixed but varies according to both object properties and the sequence of planned movements. Despite the presence of variability, this context is predictable in the sense that the worker relies on priors to estimate the required motor commands. If the worker carries out this task for a prolonged time, s/he will adjust her/his motor plan according to the object and action. In other words, motor adaptation and context switching will occur.

Robot-based investigations highlighted some limitations of the brain to concurrently learn different task dynamics (Gandolfo et al., 1996a; Conditt et al., 1997; Mussa-ivaldi, 2002), even when the expected dynamics was made fully predictable through the use of explicit cues (Krakauer et al., 1999; Osu et al., 2004). Some other contexts, however, allow the motor system to learn different dynamics if these are associated with distinct tools (Kluzik et al., 2008), objects (Ahmed et al., 2008), control policies (White and Diedrichsen, 2013) or effectors (Nozaki et al., 2006). Whether subjects can or cannot switch between contexts critically depends on experimental details.

Quite surprisingly, many examples of successful switching were also shown between altered gravity environments despite the fact these environments affect the human body in its entirety including many physiological parameters. Parabolic flights and human centrifuges provide unique means to alter the gravitoinertial environment. In the former, the participant is immersed into a repeated gravitational profile (e.g. 1g, 1.8g, 0g, 1.8g and back to 1g, where 1g is Earth gravity). Similarly, long arm human centrifuges allow programming an arbitrary gravitoinertial environment (e.g. staircase function from 1g to 3g). In contrast to conventional robotic experiments, where only the hand is perturbed, parabolic flights and rotating-room environments plunge the subject into an unexplored setting. Nearly perfect and surprisingly quick adaptation of motor responses in those challenging environments were nevertheless observed in dexterous manipulation (Hermsdörfer et al., 1999; Nowak et al., 2000; Augurelle et al., 2003; White et al., 2005; Göbel et al., 2006; Mierau et al., 2008; Crevecoeur et al., 2009; Barbiero et al., 2017) and arm movements tasks (Papaxanthis et al., 1998; White et al., 2008).

A question arises as to why switching is facilitated in radically new contexts whilst it is much more difficult in some laboratory robot-based experiments? Adaptation is a hallmark of successful tuning of internal models. In other words, our brain develops strategies to anticipate and counteract expected perturbations. To do so, it needs information and time. Visual inflows provide key information to refine our priors about an upcoming action. For instance, before lifting an object, our brain analyses different features such as size (Gordon et al., 1991a, 1991b), shape (Jenmalm and Johansson, 1997) and weight distribution (Johansson et al., 1999). All these factors influence predictive scaling of fingertip forces in dextrous manipulation. The lack of grip force adjustment is observed either through accidental slips or abnormally high safety margins. One such situation can be generated by the well-known size-weight illusion paradigm used in cognitive psychology. When participants were asked to lift a large and a small object, which seemed to be of the same material but were designed to have equal weight, peak grip and load force rates were initially scaled to object size, whereas after four trials, these signals were similar for the two objects and appropriately scaled to object weight (Flanagan and Beltzner, 2000). Therefore, after only a few practice trials, the central nervous system is capable to build two representations that can be selected on a trial basis upon context. Whether this is the same internal representation but parameterized by external information or two hard coded independent internal models remains controversial (Wolpert and Kawato, 1998).

While the importance of vision is not disputed, other sensory information, such as haptics, are processed to refine representation of internal models underlying object manipulation. We hypothesize that however radical and new the environment is, coherent sensory inflows (e.g., proprioception, vestibular) will provide much more useful information to the brain to optimize the behaviour. Here, we test the ability of seven participants to adapt to and switch between very unusual dynamical contexts generated by rotation of a long-arm human centrifuge. We measured adaptation through the robust paradigm of grip force adjustments to load force (Westling and Johansson, 1984; Jaric et al., 2005). We speculate that the nature of the environment itself will allow participants to switch between these different contexts.

## Materials and Methods

### Subjects

Seven right handed male subjects (42.1 years old, SD=9.3) participated in this experiment. A medical flight doctor checked their health status before the experiment. The protocol was reviewed and approved by the Facility Engineer from the Swedish Defence Material Administration (FMV) and an independent medical officer. The experiment was overseen by a qualified medical officer. The study was conducted in accordance with the Declaration of Helsinki (1964). All subjects gave informed consent prior to the study.

### Centrifuge facility

Centrifugation took place at QinetiQ’s Flight Physiological Centre in Linköping, Sweden. The centrifuge has a controllable swinging gondola at the end of a 9.1-meter long arm (Fig. 1A and see Levin & Kiefer (2002) for technical details). Pre-programmed G-profiles could be specified and the closed loop control of the gondola ensured that the gravitoinertial force was always aligned with body axis (Gz). Subjects were strapped while seated and cushioning was provided for comfort. Their electrocardiogram was continuously monitored during the entire centrifuge run for safety reasons. One-way video and two-way audio contacts with the control room were available at all time. In order to minimize nauseogenic tumbling sensations during acceleration and deceleration, subjects were instructed to avoid head movements. Furthermore, G-transitions between stable phases were operated below 0.32 g/s until the desired level was reached.

**Figure 1.**
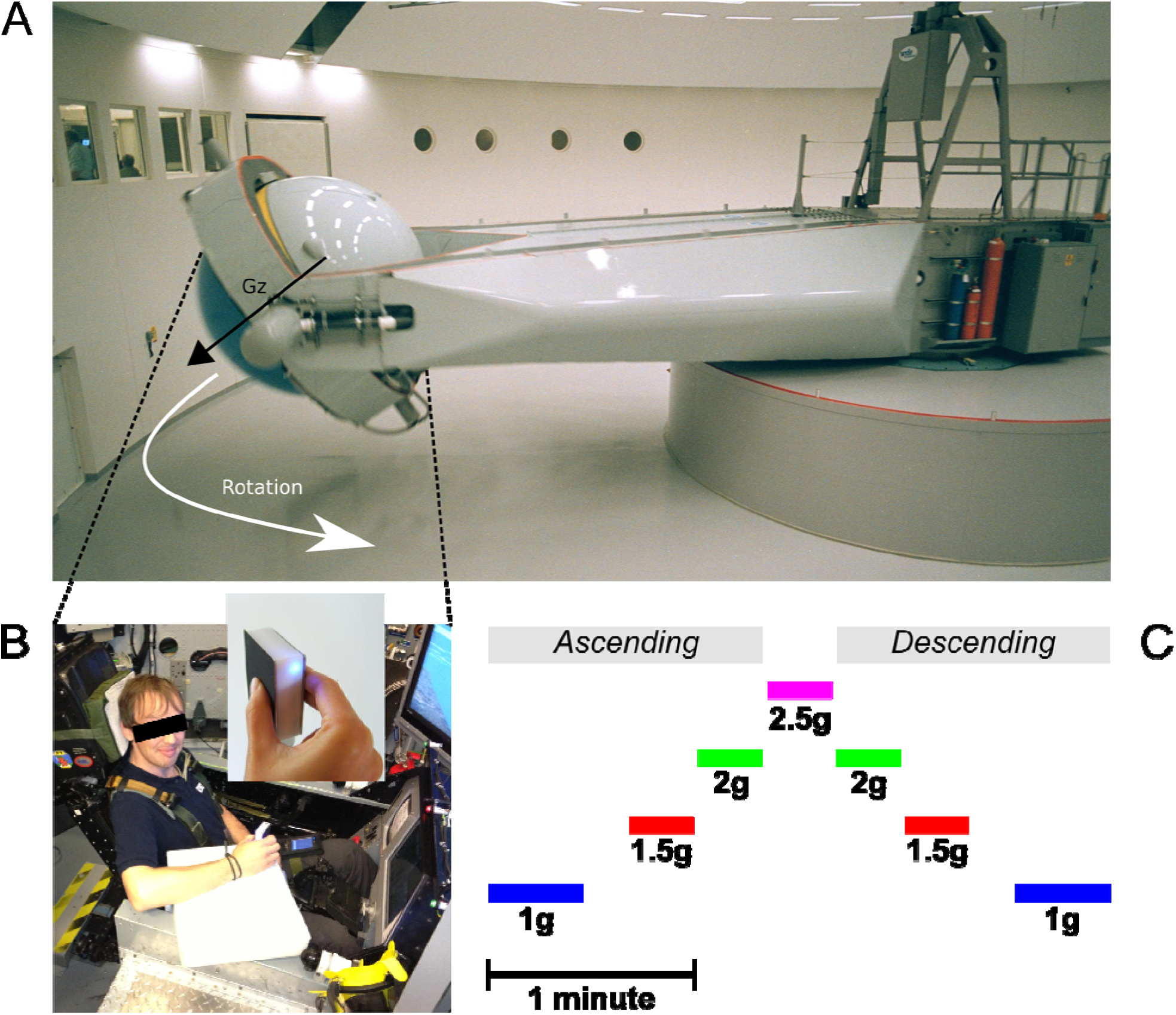
**(A)** Centrifuge room and the long-radius (9.1m) Human Centrifuge operated by QinetiQ. The control room can be seen through the windows on the left. The vector Gz illustrates the direction of the gravitoinertial force. **(B)** Subject seated in the training insert cockpit mock-up in the gondola. The manipulandum (see enlargement) was held in the right hand and was connected to a palm pilot device through a Bluetooth connection. The manipulandum was attached to the wrist and the palm device was strapped to the left thigh. A piece of white foam supported the forearm and ensured the manipulandum was in the same horizontal position between trials. **(C)** Time scaled and chronological illustration of the Gz profiles programmed in the centrifuge.

The wireless test object (mass=0.13kg) incorporated a force sensor, which measured the grip force applied against the grip surfaces (see Fig. 1B). The design of the sensor guaranteed an accuracy <±0.1N, even if the location of the center of force application was off-axis. The load force was measured along the axis of a stand until object lift-off with the same accuracy. An accelerometer that measured combined gravitational and kinematic accelerations along the object’s long axis was mounted inside the test object (accuracy ±0.2 m/s^2^). After lift-off, load force was calculated from the acceleration signal and the object mass. All signals were A/D-converted and sampled at a frequency of 120Hz. The digitized signals were then transmitted to a Palm device through a Bluetooth connection. Data were downloaded after the recordings to a standard PC for analysis.

### Procedure: lift task during centrifugation

The centrifuge was programmed to deliver a ramp up/ramp down Gz-profile for 180s (Fig. 1C). The initial 1g phases (idle) lasted for 27.4s. Then, the system was controlled to generate 1.5g, 2g, 2.5g, 2g and 1.5g. Each phase lasted 18.4s and transitions lasted 1.6s (0.31g/s). After a last transition, the system reached its final 1g phase and recording stopped after another 27.4s period. Note that transitions between 1 and 1.5g were longer (13.4s, 0.04g/s) as they were more likely to induce motion sickness. Table 1 reports mean and standard deviations of accelerations recorded during each trial and in each stable phase of the centrifugation profile and shows that the environments were very stable.

**Table 1.**
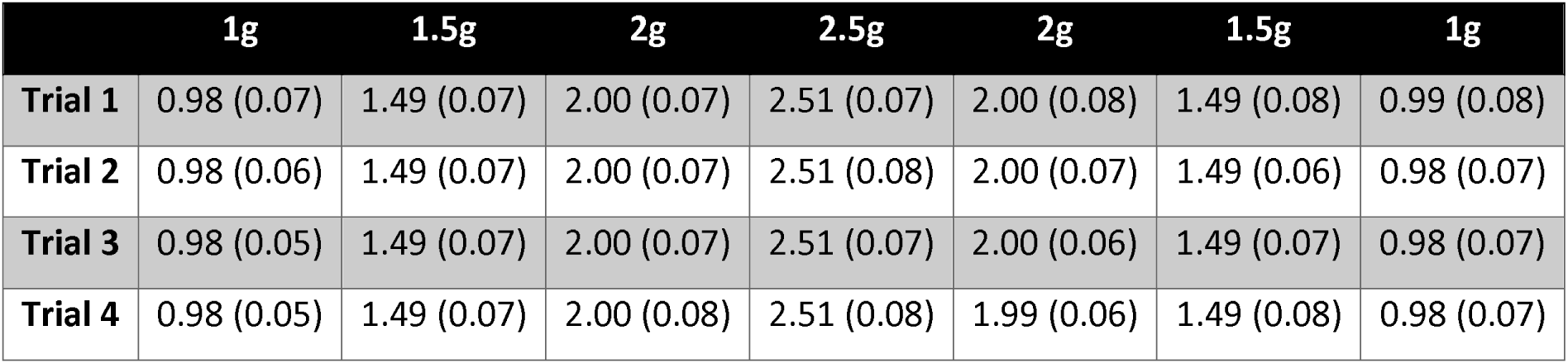
Magnitude of the local gravitoinertial acceleration (Gz) in each programmed environment in the centrifuge and during each individual lift (rows). We first calculated mean and SD of the accelerations during the trial. Cells contain the average and standard deviations of the above values across participants (N=7). The gondola rotated at a very constant rate during each phase, leading to highly reliable and stable environments.

The operator was provided with feedback about real time gravity and was in continuous verbal contact with the participant. At each GO signal (“LIFT!”), the participant adopted a precision grip configuration to grasp and lift the manipulandum. When the operator announced the STOP signal (“DOWN!”) after about 2 seconds of stationary holding, the participant gently let the object down on the support. The same task has been extensively used in previous investigations (Westling and Johansson, 1984). Four trials were completed during each stable gravitational phase. Between consecutive trials and during Gz-transitions until the first trial in the new environment, participants were asked to adopt a relaxed posture with the hand and forearm resting on the ulnar edge, and the index finger and thumb positioned approximately 2cm apart from the manipulandum grip surfaces.

### Data analysis

Grip force, load force and object acceleration along the vertical axis were low-pass filtered at 20Hz with a zero phase lag autoregressive filter. The derivatives of the force signals were then computed with a finite difference algorithm.

Figure 2 presents load force (red trace) and grip force (blue trace) in a typical trial in 1.5g that resembled those in earlier studies (Johansson and Westling, 1984; Westling and Johansson, 1984). We first determined peaks of grip force and load force for further analysis (GF_MAX_, LF_MAX_, Fig. 2). Grip force and load force onsets were identified when force rates exceeded 0.4N/s for 125ms (respectively tGF_o_ and tLF_o_). We identified the time at which load force rate fell below -2N/s for at least 125ms and subtracted 250ms to define the end of the trial. Load force and grip force plateaus were measured as the average load force and grip force during the last second of the trial. We then fitted an exponential function to the grip force profile between its peak and the end of the trial: *GF*(t) = *a* + *be*^−*ct*^ (Fig. 2). This allowed us to reliably quantify grip force decay through parameter *c* and the plateau phase of grip force with the offset parameter *a*. There was a good correlation between this last parameter and the average grip force during the last second of the trial, r=0.83, p<0.001.

**Figure 2.**
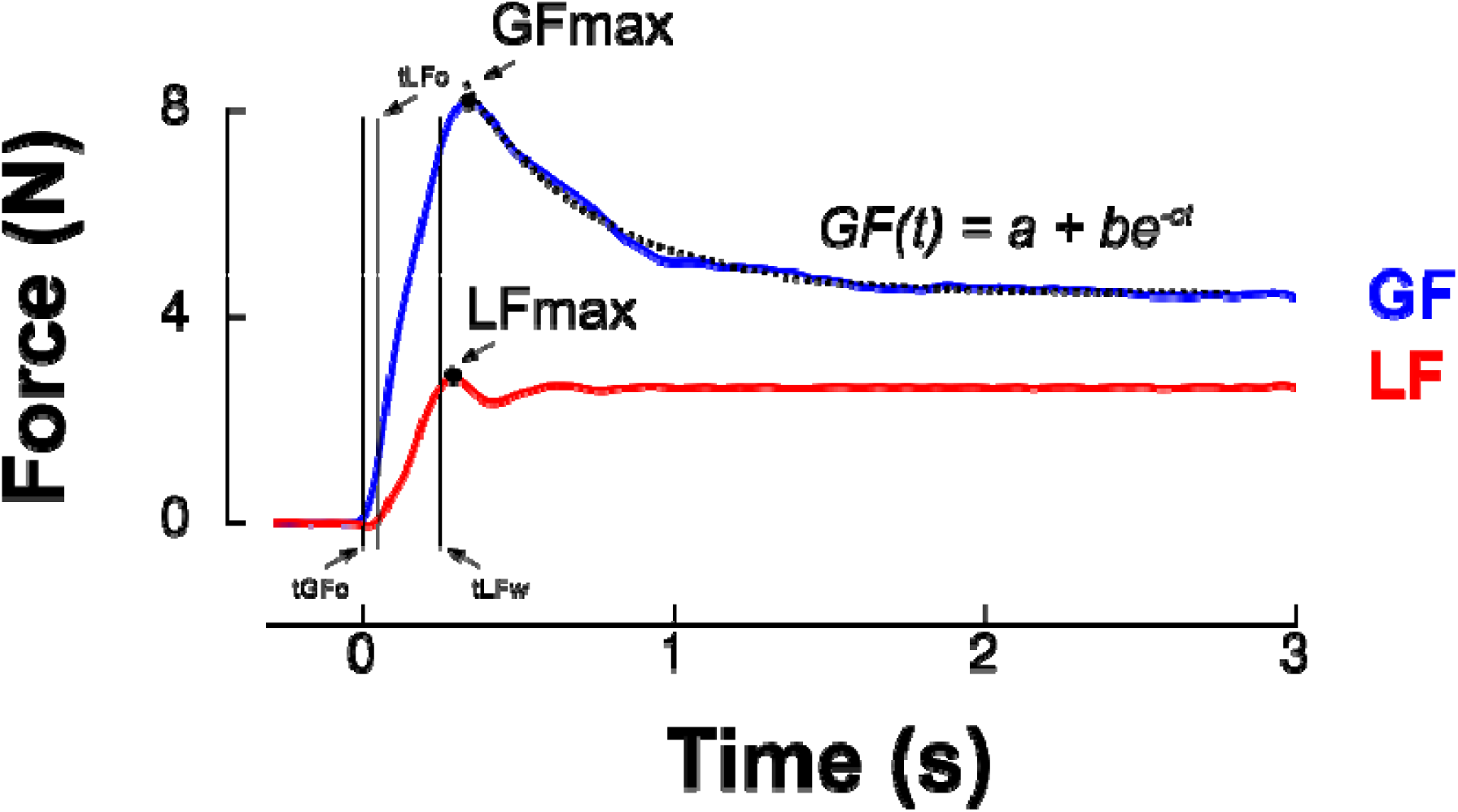
Grip force (blue trace) and load force (red trace) over time for a single lift trial. The first two vertical cursors (tGFo and tLFo) enclose the preload phase. Time 0ms corresponds to grip force onset (tGFo). The loading phase is the time between tLFo and tLFw (two last vertical cursors, see Methods). The dashed line is the best exponential fit to grip force between its peak (GFmax, black dot) and the end of the trial.Parameter a provides a more reliable estimate of grip force reached in the plateau phase than a classical average during the last portion of the trial. The rate of decrease of grip force following its maximum was quantified by parameter c.

Furthermore, two temporal parameters that characterize the grip-lift task were extracted (Fig. 2): the duration of the preload phase (delay between grip force and load force onsets, *tLF*_*o*_ – *tGF*_*o*_) and duration of the loading phase (delay between load force onset and the moment load force equals the object’s weight, *tlF*_*w*_ – *tlF*_*o*_). Finally, we calculated the cross correlation between load force rate (reference signal) and grip force rate. We shifted grip force rate between tLFo-150ms and tLFo+150ms with respect to load force rate. This procedure yielded the largest coefficient of correlation and the time shift for which this condition was fulfilled. These two values were computed for each individual trial and provided an estimate of the overall synergy of the grip-lift movement. Correlations quantified how well grip and load force profiles matched, which indicated the quality of anticipatory scaling of grip force to load force. Time-shifts provided a measure of the asynchrony between the two forces. A positive time-shift indicates that grip force led load force, as it is usually reported in healthy humans in dextrous tasks (Johansson and Westling, 1988; Forssberg et al., 1991).

Quantile-quantile plots were used to assess normality of the data. Repeated-measure ANOVAs were performed on the above variables to test for the effects of gravity (factor GRAVITY=1g, 1.5g, 2g or 2.5g) and trial (factor TRIAL=T1, T2, T3 or T4). In complementary analyses, we compared the first, ascending, 1g, 1.5g and 2g phases with the second, descending, 2g, 1.5g and 1g phases (factor PHASE=ascending or descending). Participants were only faced once to the 2.5g-phase. Therefore, it was not included in the ANOVA when factor PHASE was considered. Post hoc Scheffé tests were used for multiple comparisons and paired t-test of individual subject means were used to investigate differences between conditions. Alpha level was set at 0.05. Because the sample size is moderate (n=7 participants), partial eta-squared are reported for significant results to provide indication on effect sizes. The dataset was visually inspected to ensure these parameters were accurately extracted by custom routines developed in Matlab (The Mathworks, Chicago, IL).

## Results

Participants performed a precision grip lifting task when the gravitoinertial environment was varied with a centrifuge. Figure 3A depicts average load force during the plateau phase in each gravitational environment separately for each trial. Since the object was held stationary during that period, the load force reflects the weight of the manipulandum during the respective G-levels. Consistently, a 3-way RM ANOVA shows that load force plateau – or object weight – was only influenced by GRAVITY (F(2,131)=143920.04, p<0.001, 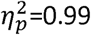) and not TRIAL (F(3,131)=0.8, p=0.494) or PHASE (F(1,131)=0.3,p=0.619), with no interaction effect (all F<1.6, all p>0.191). Participants matched this level of static load force with grip force, as illustrated in Figure 3B. Again, the ANOVA only reported a main effect of GRAVITY, F(3,131)=28.9, p<0.001, 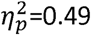, with no significant TRIAL, PHASE or interaction effect (all F<2.9, p>0.094). The ratio between grip force and load force was not affected by any of the factors (Fig. 3C; all F<0.4, all p>0.318) except PHASE (F(1,131)=5.0, p=0.025, 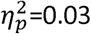). We indeed found a small decrease of the safety margin during the plateau phase in the second (descending) phase compared to the first (ascending) phase (ascending: 1.55; descending: 1.37).

**Figure 3.**
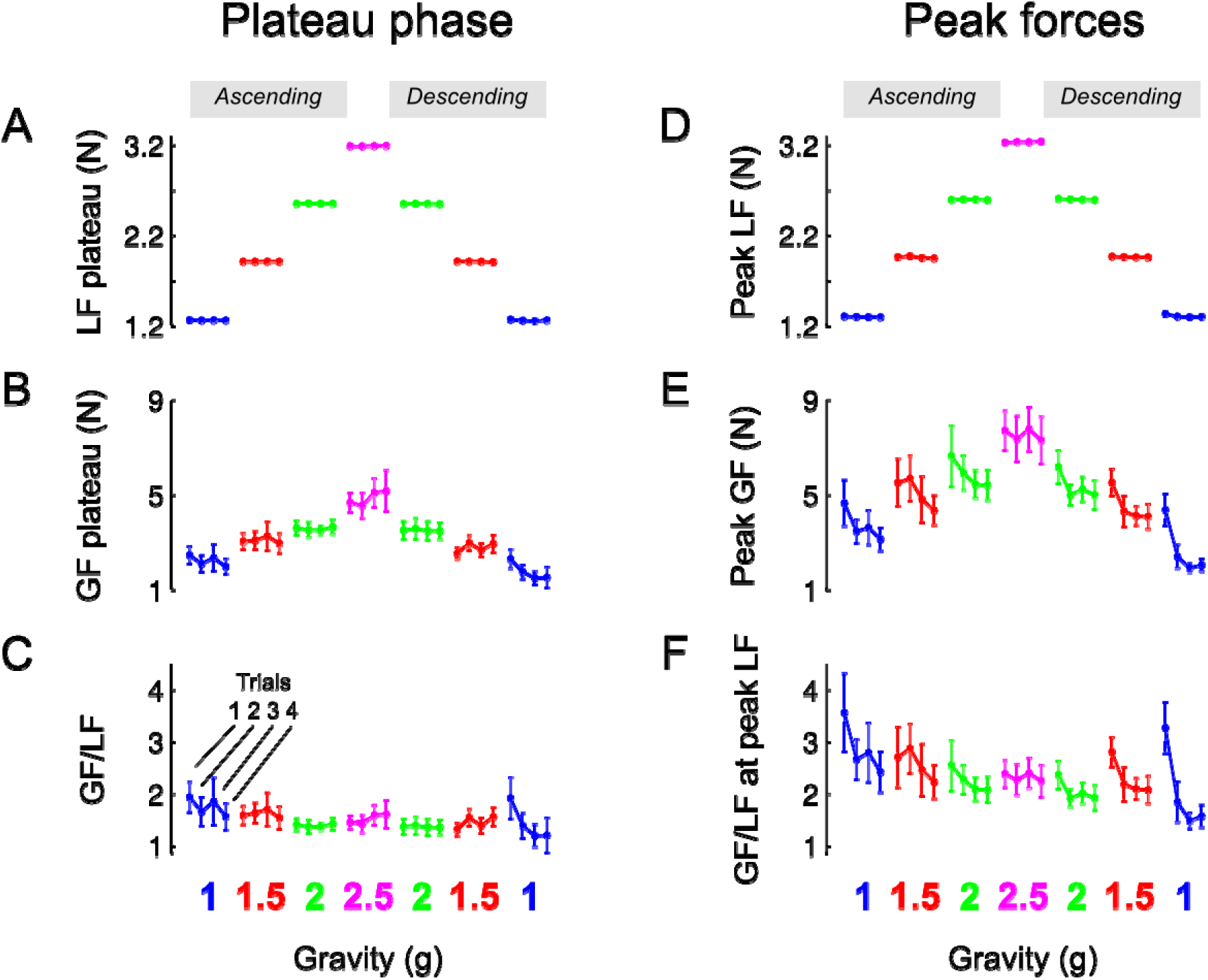
Mean and SEM of parameters of the task that characterize the plateau phase (left column, **A-C**) and when load force reached a maximum (right column, **D-F**). Data are presented in chronological order, following the successive exposures to 1g, 1.5g, 2g, 2.5g, 2g, 1.5g and back to 1g (same colour code as in Fig. 1). Data are also shown separately for each trial in a given environment (see panel C, 1g). The ‘ascending’ (resp. ‘descending’) phase comprised the increasing (resp. decreasing) gravitoinertial environments 1g to 2.5g (resp. 2.5g to 1g).

Load force depends on gravity (mass x gravitational acceleration) and kinematics through its inertial component (mass x acceleration). Peaks of load force were on average only 12.3% larger than load force plateau (compare Fig. 3A and D). The statistical analysis again reported a main GRAVITY effect on peak load force (Fig. 3D, F(3,131)=2769.2, p<0.001, 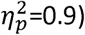). Furthermore, load force peaks were also 8% smaller in the descending phase PHASE (PHASE: F(1,131)=7.4, p=0.007, 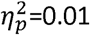). Participants moved the manipulandum in such a way that apart from the somewhat lower acceleration during decent, it was not influenced by any other factor (all F<0.9, p>0.495).

Figure 3E shows that grip force peaks were adjusted to load force (main effect of GRAVITY, F(2,131)=18.8, p<0.001, 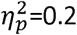). Interestingly, Figure 3E also shows two additional effects. On the one hand, grip force peaks decreased across trials during the first and second exposures to 1g, 1.5g and 2g. On the other hand, the average level of peak grip force seemed to be lower in the second, descending, phase. An ANOVA confirmed that grip force peaks decreased with TRIAL (F(3,131)=4.7, p=0.004, 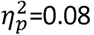) and were lower in the descending phase of the profile (PHASE, F(1,131)=5.1, p=0.025, 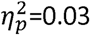), without any interaction (all F<0.5, all p>0.643). Post hoc tests revealed that trial 1 was marginally different from trials 2-4 (p<0.045). When the ANOVA was conducted only with trials 2, 3 and 4, it yielded no results, F(2,97)=0.7, p=0.496. This effect is also reflected in the ratio between grip and load forces when load force was maximum (Fig. 3F; TRIAL: F(3,131)=4.8, p=0.003, 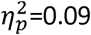 and PHASE: F(1,131)=7.1, p=0.009, 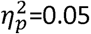). Finally, we also found that, within trials, grip force decreased faster to its value in the plateau in the descending phase (parameter c, exponential decay in ascending vs. descending: 714ms vs 384ms, main effect of PHASE F(1,131)=4.6, p=0.035, 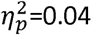) with no other effects (all F<0.73, all p>0.588). Altogether, these results show that participants adopted stereotyped movements from the very first trial in every gravitational environment and decreased grip force across trials and exposure according to the g-level.

The grip force level during the plateau phase was adjusted more than a second after first contact occurred with the object and this regulation was probably influenced by feedback mechanisms. Although grip force peaks occurred rather early after lift-off (mean across participants=273.2ms, SD=58.8ms), peak grip force rates, which always occur earlier than grip force peaks (mean=104.2ms, SD=37.5ms) are therefore sometimes considered a reliable measure of feedforward processes. Interestingly, the same analysis as above led to even more significant conclusions: Peak grip force rates were proportional to GRAVITY (F(3,131)=17.0, p<0.001, 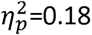), decreased with TRIAL (F(3,131)=4.2, p=0.007, 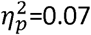) and pwere lower in the descending phase of the profile (PHASE, F(1,131)=5.1, p=0.025, 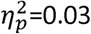) with no interaction (all F<0.5, all p>0.712).

This anticipatory strategy is compatible with the very short lags observed between load force rate and grip force rate (mean=1.7ms, SD=7.3ms) as well as associated high correlations (mean=0.91,SD=0.1). A two-way ANOVA revealed that the correlation increased with TRIAL (Fig. 4A, F(3,93)=4.2, p=0.008,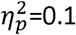), was not affected by GRAVITY (F(3,93)=1.69, p=0.17) and that the lag was not significantly altered (Fig. 4B, all F<0.51, all p>0.123). The ANOVA did not report any significant result when we excluded trial 1, F(2,65)=0.8, p=0.452.

**Figure 4.**
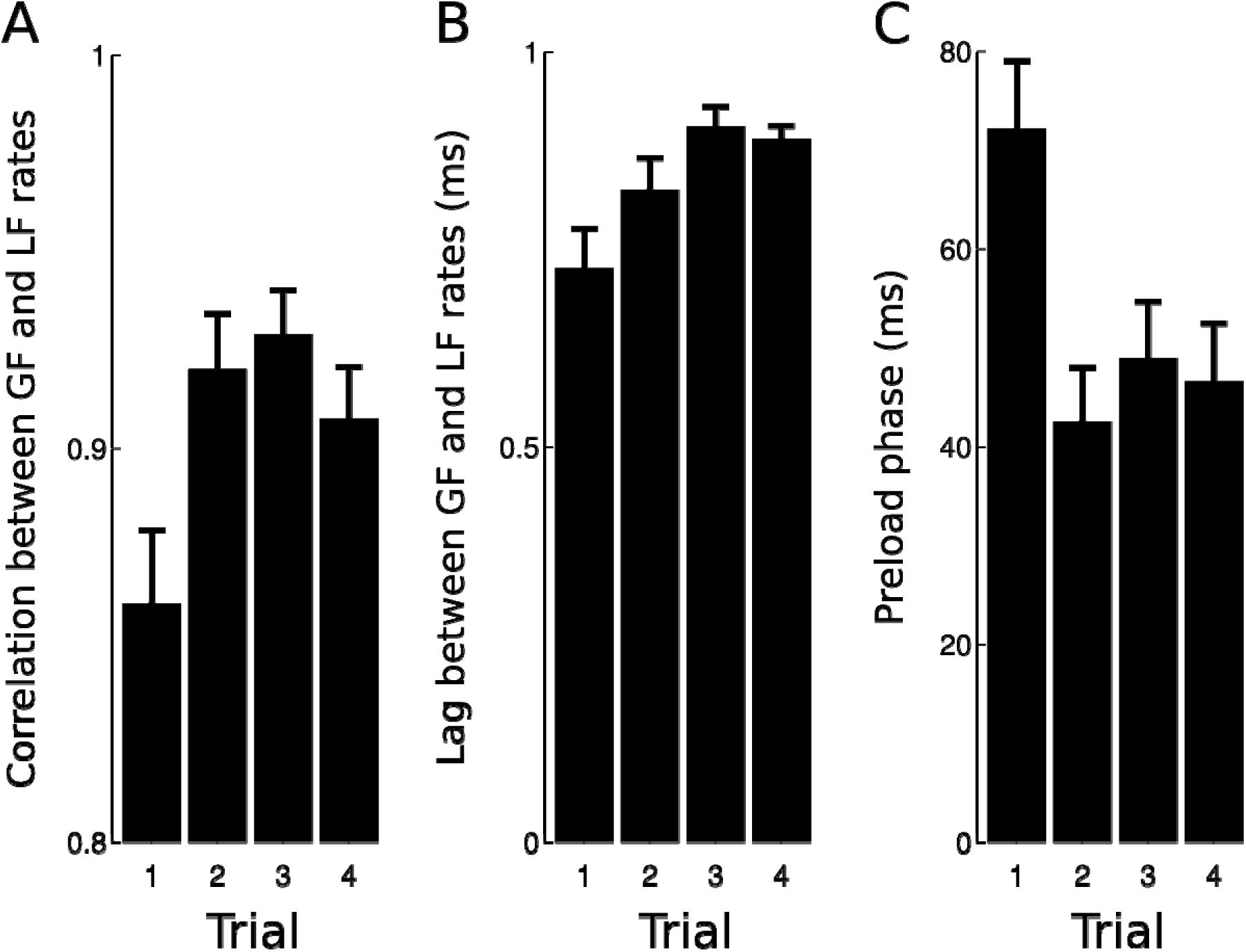
Effects of trial number on the correlation **(A)** and lag **(B)** calculated between load force rate and grip force rate during the loading phase. Trial 1 had a lower correlation than trials 2-4. **(C)** The preload phase was the longest during trial 1 in comparison with trials 2-4.

The preload phase, i.e. the delay between object-finger(s) contact and the first increase in load force, is an important underlying variable that characterizes a grip-lift task because, during this short period of time, physical properties of the grasped object are encoded by mechanoreceptors. Figure 4C depicts average preload phases in the four trials. The ANOVA reported a significant effect of TRIAL, F(3,93)=3.8, p=0.013, 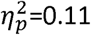, with no other effect (all F<0.94, all p>0.423). A post hoc test showed that the preload phase in trial 1 was longer than during trials 2-4 (70.9ms to 46.9ms, 33.8% drop, p=0.041).

The most striking finding is that subjects could adjust grip force to load force from the very first trial in the new environment. To quantify this ability, we calculated the correlation between load force and grip force peaks in each phase but only for trial 1. We found very good and similar correlations in the ascending and descending phases (Fig. 5). When we pooled phases together, the correlation reached r=0.91 and was significant (p<0.001).

**Figure 5.**
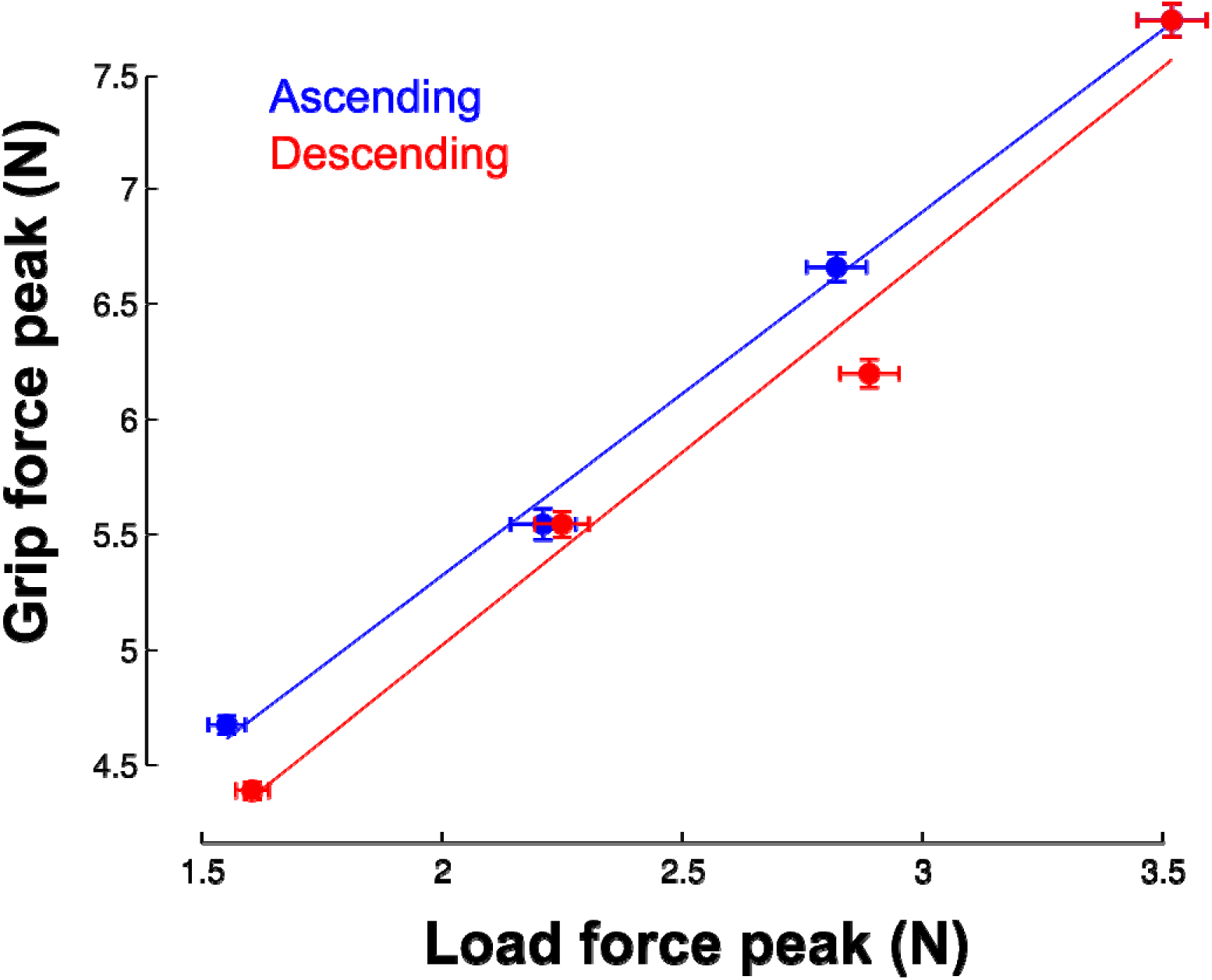
Correlation between load force peaks (x-axis) and grip force peaks (vertical axis) in the first trial and separately for each phase. Each point corresponds to the average across the seven participants in each of the four gravitoinertial contexts. The linear regressions were significant in both ascending (r=0.99, p=0.002, slope=1.6, offset=2.2) and descending phases (r=0.99, p=0.012, slope=1.67, offset=1.68) Vertical and horizontal error bars correspond to STD. Note that the point in the upper right corner was identical in the ascending and descending phases (only one 2.5g phase).

In the previous sections, we showed that although subjects moved the object consistently across conditions, grip force was not completely adapted upon entry in the new environment. Indeed, there were genuine differences between trial 1 and the three following trials in the same condition. Figure 6 illustrates average load force (red) and grip force (blue) profiles in the first trial (T1, solid line) and in the last trial (T4, dashed line). The upper row reports these time series in the ascending phase (Fig. 6A-D, 1g, 1.5g, 2g and 2.5g) and the lower row depicts these data in the descending phase (Fig. 6E-H, 2.5g, 2g, 1.5g and 1g). Note that for the sake of clarity and comparison, panels D and E report data from the same grip and load force profiles in 2.5g. While Figure 6 shows that load forces overlapped between trial 1 and trial 4, grip force was always larger in trial 1 compared to trial 4.

**Figure 6.**
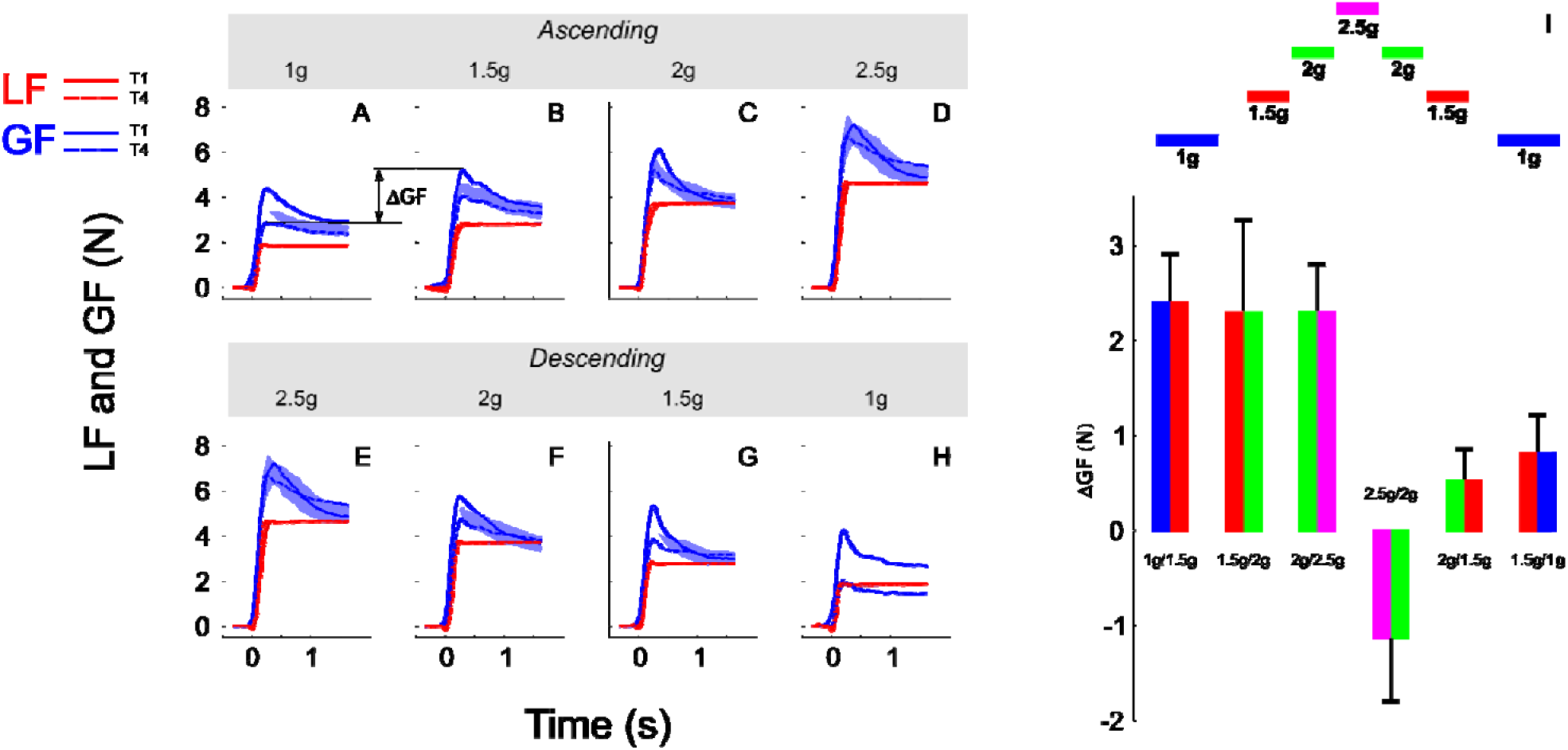
**(A-H)** Grip force (blue traces) and load force (red traces) over time plotted in each gravitoinertial environment and for the first (T1, solid lines) and last (T4, dashed lines) trial. Traces were the averages across 7 participants and the shaded area corresponds to SEM. Notice that load force overlapped closely between T1 and T4 in all conditions. The upper row **(A-D)** corresponds to the ascending phase and th lower row depicts time series in the descending phase **(E-H)**. For clarity, since we had only one 2.5g environment, panels D and E present the same data. The index ΔGF (illustrated in panel A) quantifies the switching between environments and is calculated as the first grip force peak in the next environment (at trial 1) minus the last grip force peak reached in the current environment (at trial 4). **(I)** Average and SEM of ΔGF for each of the six transitions. Bar plots are bicolour; left colour corresponds to the current environment and right colour corresponds to the next environment (refer to the sketch above).

We quantified the participants’ ability to switch between environments by subtracting grip force peak recorded in the last trial in the previous gravity level from grip force peak during the first trial in the next environment, 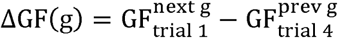. This index, ΔGF, is illustrated between 1g and 1.5g in the ascending phase in Figure 6 between panels A and B. We ran an ANOVA with factors PHASE (ascending vs. descending) and a new factor that characterizes the two environments between which ΔGF is calculated (SWITCH: 1g to 1.5g, 1.5g to 2g and 2g to 2.5g). The ANOVA reported that ΔGF was significantly larger in the ascending phase (F(1,34)=19.05, p<0.001, _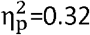_). These effects are illustrated in Figure 6I (see also Table 2). Similar results were found when peak grip force rates were used to calculate the index. Furthermore, a t-test showed that ΔGF was significantly larger than 0 in the ascending phase (mean=2.33N; t(20)=6.05, p<0.001, 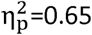), but not in the descending phase (mean=-0.01N; t(18)=-0.02, p=0.984). This analysis reveals an asymmetric behaviour between phases, suggesting that the peak grip force in the current environment is not only planned on the basis of the performance in the last trial in the previous environment combined with the anticipated effects of the upcoming gravitoinertial context. As outlined below, a strategy consisting in adopting a safety margin that is large in the first trial in a new environment but dissipates in the following trials could be relevant.

**Table 2.** Values of different parameters between two consecutive trials in two different environments (Trials). Are reported: step of LF and GF and the predictive (*αm*Δ*g*) and uncertainty (*β*) terms under two different hypotheses (“Prediction correct” and “Prediction incorrect”). The bold italic row highlights the first descending step, i.e., between 2.5g and 2g.

| Trials | $\Delta LF$ (N) | $\Delta GF$ (N) | Correct Prediction | | Incorrect Prediction | | |
| --- | --- | --- | --- | --- | --- | --- | --- |
| | | | $\alpha m\Delta g$ (N) | $\beta$ (N) | $\alpha m\Delta g$ (N) | $\beta$ (N) | $\alpha$ |
| **1 to 1.5** | 0.70 | 2.39 | 0.96 | 1.44 | 0.95 | 1.44 | 1.50 |
| **1.5 to 2** | 0.61 | 2.29 | 0.96 | 1.33 | 0.85 | 1.44 | 1.33 |
| **2 to 2.5** | 0.64 | 2.30 | 0.96 | 1.34 | 0.86 | 1.44 | 1.35 |
| **2.5 to 2** | **-0.60** | **-1.12** | **-0.96** | **-0.16** | **-2.56** | **1.44** | **4.01** |
| **2 to 1.5** | -0.60 | 0.52 | -0.96 | 1.48 | -0.92 | 1.44 | 1.44 |
| **1.5 to 1** | -0.66 | 0.81 | -0.96 | 1.77 | -0.63 | 1.44 | 0.99 |

A question naturally arises as to why this ΔGF is asymmetric while step of load forces are symmetric between phases? The ANOVA reported that ΔLF, which equals *mg*_*t*+1_ – *mg_t_*, calculated between peaks of load force, were of course different between the ascending and descending phases (F(1,34)=15.6, p<0.001, 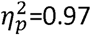) but did not vary within phase (F(2,34)=0.43, p=0.654) and were symmetric (t(18)=-1.4, p=0.167). Table 2 reports these values of ΔGF. The prediction of what increment of grip force to apply can be based on the expected increment of load force. These forces have been shown to be reliably linearly correlated with a gain *α*: Δ*GF* = *αΔLF* = *α*(*LF*_*t*+1_ - *LF_t_*), where t denotes the last trial in the previous context and t+1 the first lift in the next context. We assume that similar accelerations were produced by participants, which yields Δ*GF* = *α*(*mg*_*t*+1_ – *mg_t_*) = *αmΔg*. Furthermore, participants often adopt some security margin *β* that reflects task and environmental uncertainties and risk aversion. The increment of grip force can therefore follow this simple rule: Δ*GF* = *αm*Δ*g* + β, where the first term quantifies prediction based on experienced and expected information and the last term includes uncertainty.

We can now test two alternative hypotheses to explain why ΔGF does not follow the simple model described above (Fig. 6I). On the one hand, the prediction can be correct and constant but uncertainty can vary. We set the value of the gain *β*; to 1.5, which corresponds to the mean of the grip to load force ratio in all first trials in a new environment (Fig. 3C). Table 2 (correct prediction, *αmΔg*) reports values of the predictive term that are proportional to ΔLF. In order to match the observed ΔGF, the second term had to be adapted (Table 2, correct prediction, β). Alternatively, if we set an uncertainty value to the constant 1.44N that corresponds to uncertainty measured in a normal case (1g), the term αmΔg becomes variable, which is caught in α (Table 2, α). In the first hypothesis, uncertainty is rather constant in the ascending phase but jumps to a negative value before increasing again in the descending phase. Usually, uncertainty decreases over time, when one gets more confidence in the task. Instead, our data seem to favour the second hypothesis. In that case, the internal model is wrongly adjusted, especially in the first descending step (Table 2, bold and italic row).

## Discussion

Humans use many different objects in various situations. For instance, when cooking, one hand can move an egg off the table and shortly after, manipulate a heavy pan. Fortunately, the brain developed strategies that allow to anticipate task-relevant parameters and adjust the control policy accordingly, from the very first instant we start the task. This ability has been demonstrated in the past in several experimental contexts and is formalized by the concept of internal models. In particular, it was suggested that the brain can store multiple predictive models and select the most appropriate one according to the task at hand (Wolpert and Kawato, 1998).

Importantly, these models are flexible. Most of the time, participants can learn the appropriate dynamics of a new task in a matter of a few trials. Quite surprisingly, this also holds when the environment is radically altered like in parabolic flight (Nowak et al., 2000; Augurelle et al., 2003) or when subjects are confronted to artificial new dynamics (Flanagan and Wing, 1997). It seems that, after sufficient training, participants can switch between these models effortlessly. However, force-field learning experiments show that two dynamic internal models cannot be learned concurrently unless the posture of the arm is changed between conditions (Gandolfo et al., 1996b; Mussa-ivaldi, 2002). Interestingly, abstract representations of different objects can be combined in the brain to create a new one, adapted to a new situation. In a nicely designed paradigm, participants trained to lift objects of different masses and were then asked to lift the combined object (Davidson and Wolpert, 2004). Grip force rates were adjusted predictively in the very first trial for the combined object, suggesting they stacked both previously formed models. How do we reconcile experimental contexts in which adaptation needs time and others that do not, or, in other words, that allow switching? We posit that a fundamental difference between these conditions is the availability of different sensory information that allow much more efficient adaptation.

Here, we asked participants to lift a lightweight object but in different gravitoinertial environments generated by a long arm human centrifuge. The dynamics of the system and the visual environment in the gondola were such that the different gravitoinertial levels were felt like pure gravitational increments. Subjects are extremely familiar with the employed lifting task and with this kind of object but not at all with the environment. Before the experiment, they were told what gravitoinertial profile (amplitude and time course) was implemented in the system. They were also warned in real time, during the experiment, when a new transition was about to occur. All participants had therefore a cognitive knowledge but not (yet) a multisensory experience of the task.

We found a remarkable ability of participants to regulate their grip force from the outset. How was that possible? First, the brain could use information from all sensory modalities. This task, once the object was contacted by the fingers, relied mostly on tactile and proprioceptive feedback. Initial perfect adaptation underlines the importance of that sensory modality. This is in agreement with the work mentioned above (Davidson and Wolpert, 2004) since in that study, participants were prevented from any visual or auditory cue. However, Davidson and Wolpert’s experiment was conducted in a familiar, terrestrial, environment. Second, participants also had a theoretical knowledge of the environment. However, it was also shown that pure cognitive knowledge about a change of context is sometimes insufficient to allow prediction. For instance, when subjects decreased the weight of a hand-held glass of water by drinking with a straw, they could match the change of weight with grip force which they couldn’t when lifting the object after drinking while the object was left on the table (Nowak and Hermsdörfer, 2003). Similarly, the prediction of the effects of gravity of a falling virtual object was only possible when a physical interaction with the object was required (Zago et al., 2004). Third, repeating the same trial many times triggers use-dependent mechanisms (Diedrichsen et al., 2010). This propensity of performing the same action if it was successful during the previous trials is responsible for the appearance of large errors if a contextual parameter is changed unbeknownst to the participant. While this process may have been used within a gravitoinertial phase, it was certainly not the case between phases. Altogether, this suggests that multisensory information is essential to switch between environments. Two learning mechanisms may both contribute to adaptation during this task but their respective importance may be weighted differently. Prediction errors are used by error-based learning processes when switching while use-dependent mechanisms are active within each constant environment.

Despite the fact we observed good overall adaptation of grip force to load force in all phases, there were nevertheless exceptions noticeable at two different timescales. On the one hand, when comparing equivalent environments, grip forces were smaller in the second, descending, phase of the experiment. This is also reflected by a faster decay of grip force to a smaller plateau value. During a parabolic flight campaign, the static grip force produced to hold an object stationary was massively increased during the first experience of 0g and 1.8g suggesting a strong effect of stress induced by the novel environmental conditions (Hermsdörfer et al., 1999). This increase in grip force levels resolved however quickly across the subsequent exposures to the new gravitoinertial conditions. This behaviour may reflect habituation and not a change in motor prediction. On the other hand, there were subtle adjustments in grip force (not load force) between the first trial and the next trials. Namely, peak grip force, grip force rate, the grip to load ratio and the preloading phase all decreased after the first trial and the synergy between both forces improved. Therefore, a pure, perfect switch really needs one trial to occur.

It is immediately clear in Figure 6 that the change in grip force directly after a change of g-level does not directly reflect the change in load. It rather seems that the change in grip force is exaggerated since it is reduced substantially in the following trials. In all environments, a safety margin, linked to self-perception of uncertainty, was obviously employed during the first contact with the object in the new gravitoinertial environment. This margin decreased with time and confidence. The only exceptions are the trials in the highest g-level, 2.5 g. In that extreme situation, subjects experienced the highest mental and physical stress and may not have relaxed during the duration of the 2.5-g interval.

Interestingly, the shape of the grip force switch was asymmetric between ascending and descending g-changes. One reason may be that the second (descending) phase was not entirely novel for participant. This is particularly true for the transition from 2.5g down to 2g, since the ascending 2g phase was still in the recent sensorimotor history. Furthermore, because inertial fluctuations were weak, predicting the weight could have been sufficient to adjust grip force. It seems that a combination of grip force prediction according to the change in the gravitoinertial environment and a separate safety margin can predict the data quite accurately. A simple linear model that includes (1) a gain factor which reflects the calculation of the grip force change from the load change and (2) a constant magnitude of grip force increase as safety margin approximate the data well. This factor seems however not constant, but may depend on context like ascending or descending g-levels, time in the experiment, or experience with g-changes.

Finally, our data should also be put in the perspective of more theoretical motor control considerations. Despite the very new context, participants never dropped the object. In the presence of such environmental uncertainty, what strategy does the central nervous system adopt to predict a feedforward grip force command in the new phase condition? One approach consists in minimizing the squared error of potential feedforward predictions (Körding et al., 2004) i.e. penalizing too high grip forces. This can be achieved by averaging previous lifts (Scheidt et al., 2012; Hadjiosif and Smith, 2015) or using a Bayesian framework (Körding and Wolpert, 2004). The latter is more flexible as in addition to estimating physical properties linked to the object, it can also build a representation of environment uncertainty. Once both are integrated, a point estimate can be formed. By a genuine manipulation of probability distribution of object masses, a recent study showed that the sensorimotor system indeed uses a minimal squared error strategy to predict grip force (Cashaback et al., 2017). This view is not quite compatible with our results, as it does not explain the switching we observe. Another approach consists in selecting the feedforward prediction that is most likely to be correct. This strategy has been shown to occur in sequential object lifting. When confronted to lift objects of increasing weights, participant expect the next trial to be even heavier (Mawase and Karniel, 2010). Here, participants that were immersed in these gravitoinertial contexts could have formed a reliable representation of the object dynamics in the environment. This could have provided solid information in order to infer a good prediction and therefore a good switch. This is further supported by the fact grip forces were even smaller during the second descending phase Overall, this view is compatible with the fact both mechanisms are implemented in parallel (Cashaback et al., 2017). However, psychological factors such as stress, could be responsible for the asymmetry observed in the switching between ascending and descending phases. One way to address this would be to perform the same experiment as Mawase and Karniel (2010) but using decreasing weights in the laboratory environment.

## Acknowledgements

This research was supported by the European Space Agency (ESA) in the framework of the Delta-G Topical Team, the « Institut National de la Santé et de la Recherche Médicale » (INSERM) and the « Conseil Général de Bourgogne » (France).

## References

Ahmed A a, Wolpert DM, Flanagan JR. Flexible representations of dynamics are used in object manipulation. Curr Biol 18: 763–8, 2008.

Augurelle A-S, Penta M, White O, Thonnard J-L. The effects of a change in gravity on the dynamics of prehension. Exp brain Res 148: 533–40, 2003.

Barbiero M, Rousseau C, Papaxanthis C, White O. Coherent multimodal sensory information allows switching between gravitoinertial contexts. Front Physiol 8: 290, 2017.

Cashaback JGA, McGregor HR, Pun HCH, Buckingham G, Gribble PL. Does the sensorimotor system minimize prediction error or select the most likely prediction during object lifting? J Neurophysiol 117: 260–274, 2017.

Conditt MA, Gandolfo F, Mussa-Ivaldi FA. The motor system does not learn the dynamics of the arm by rote memorization of past experience. J Neurophysiol 78: 554–560, 1997.

Crevecoeur F, Thonnard JL, Lefèvre P. Forward models of inertial loads in weightlessness. Neuroscience 161: 589–98, 2009.

Davidson PR, Wolpert DM. Internal models underlying grasp can be additively combined. Exp Brain Res 155: 334–340, 2004.

Diedrichsen J, White O, Newman D, Lally N. Use-dependent and error-based learning of motor behaviors. J Neurosci 30: 5159–5166, 2010.

Flanagan JR, Beltzner MA. Independence of perceptual and sensorimotor predictions in the size-weight illusion. Nat Neurosci 3: 737–41, 2000.

Flanagan JR, Wing AM. The role of internal models in motion planning and control: evidence from grip force adjustments during movements of hand-held loads. J Neurosci 17: 1519–1528, 1997.

Forssberg H, Eliasson AC, Kinoshita H, Johansson RS, Westling G. Development of human precision grip I: Basic coordination of force. Exp Brain Res 85: 451–457, 1991.

Gandolfo F, Mussa-Ivaldi FA, Bizzi E. Motor learning by field approximation. Proc Natl Acad Sci U S A 93: 3846–3846, 1996a.

Gandolfo F, Mussa-Ivaldi FA, Bizzi E, Sciences C. Motor learning by field approximation. Proc Natl Acad Sci U S A 93: 3846–3846, 1996b.

Göbel S, Bock O, Pongratz H, Krause W. Practice ameliorates deficits of isometric force production in +3 Gz. Aviat Sp Environ Med 77: 586–591, 2006.

Gordon AM, Forssberg H, Johansson RS, Westling G. Visual size cues in the programming of manipulative forces during precision grip. Exp Brain Res 83: 482–482, 1991a.

Gordon AM, Forssberg H, Johansson RS, Westling G. Integration of sensory information during the programming of precision grip: comments on the contributions of size cues. Exp brain Res 85: 9–9, 1991b.

Hadjiosif a. M, Smith M a. Flexible Control of Safety Margins for Action Based on Environmental Variability. J Neurosci 35: 9106–9121, 2015.

Hermsdörfer J, Marquardt C, Philipp J, Zierdt A, Nowak D, Glasauer S, Mai N. Grip forces exerted against stationary held objects during gravity changes. Exp Brain Res 126: 205–214, 1999.

Jaric S, Knight CA, Collins JJ, Marwaha R. Evaluation of a method for bimanual testing coordination of hand grip and load forces under isometric conditions. J Electromyogr Kinesiol 15: 556–563, 2005.

Jenmalm P, Johansson RS. Visual and somatosensory information about object shape control manipulative fingertip forces. J Neurosci 17: 4486–4499, 1997.

Johansson RS, Backlin JL, Burstedt MKO. Control of grasp stability during pronation and supination movements. In: Experimental Brain Research. 1999, p. 20–30.

Johansson RS, Westling G. Roles of glabrous skin receptors and sensorimotor memory in automatic control of precision grip when lifting rougher or more slippery objects. Exp Brain Res 56: 550–564, 1984.

Johansson RS, Westling G. Coordinated isometric muscle commands adequately and erroneously programmed for the weight during lifting task with precision grip. Exp Brain Res 71: 59–71, 1988.

Kluzik J, Diedrichsen J, Shadmehr R, Bastian AJ. Reach adaptation: what determines whether we learn an internal model of the tool or adapt the model of our arm? J Neurophysiol 100: 1455–1464, 2008.

Körding KP, Ku SP, Wolpert DM. Bayesian estimation in force integration. J Neurophysiol 92: 3161–3165, 2004.

Körding KP, Wolpert DM. Bayesian integration in sensorimotor learning. Nature 427: 244–247, 2004.

Krakauer JW, Ghilardi M, Ghez C. Independent learning of internal models for kinematic and dynamic control of reaching. 2, 1999.

Levin B, Kiefer D. Dynamic Flight Simulator for Enhanced Pilot Training. SAFE Eur: 1–5, 2002.

Mawase F, Karniel A. Evidence for predictive control in lifting series of virtual objects. Exp Brain Res 203: 447–452, 2010.

Mierau A, Girgenrath M, Bock O. Isometric force production during changed-Gz episodes of parabolic flight. Eur J Appl Physiol 102: 313–8, 2008.

Mussa-ivaldi AKFA. Does the motor control system use multiple models and context switching to cope with a variable environment[⍰]? (2002). doi:10.1007/s00221-002-1054-4.

Nowak DA, Hermsdörfer J. Sensorimotor memory and grip force control[⍰]: does grip force anticipate a self-produced weight change when drinking with a straw from a cup[⍰]? 18, 2003.

Nowak ZD, Hermsdörfer J, Marquardt C, Philipp J, Zierdt A, Nowak D, Glasauer S, Mai N, Nowak ZD. Moving weightless objects. Grip force control during microgravity. Exp Brain Res 132: 52–64, 2000.

Nozaki D, Kurtzer I, Scott SH. Limited transfer of learning between unimanual and bimanual skills within the same limb. Nat Neurosci 9: 1364–6, 2006.

Osu R, Hirai S, Yoshioka T, Kawato M. Random presentation enables subjects to adapt to two opposing forces on the hand. 7: 111–112, 2004.

Papaxanthis C, Pozzo T, Popov KE, McIntyre J. Hand trajectories of vertical arm movements in one-G and zero-G environments. Evidence for a central representation of gravitational force. Exp Brain Res 120: 496–502, 1998.

Scheidt RA, Dingwell JB, Mussa-ivaldi FA, Vasudevan EVL, Torres-oviedo G, Morton SM, Yang JF, Amy J. Learning to Move Amid Uncertainty Learning to Move Amid Uncertainty.

Westling G, Johansson RS. Factors influencing the force control during precision grip. Exp brain Res 53: 277–84, 1984.

White O, Bleyenheuft Y, Ronsse R, Smith AM, Thonnard J-L, Lefèvre P. Altered gravity highlights central pattern generator mechanisms. J Neurophysiol 100: 2819–24, 2008.

White O, Diedrichsen J. Flexible Switching of Feedback Control Mechanisms Allows for Learning of Different Task Dynamics. PLoS One 8, 2013.

White O, McIntyre J, Augurelle AS, Thonnard JL, Mcintyre J. Do novel gravitational environments alter the grip-force/load-force coupling at the fingertips? Exp Brain Res 163: 324–334, 2005.

Wolpert DM, Kawato M. Multiple paired forward and inverse models for motor control. [Online]. Neural Netw 11: 1317–29, 1998. http://www.ncbi.nlm.nih.gov/pubmed/12662752.

Zago M, Bosco G, Maffei V, Iosa M, Ivanenko YP, Lacquaniti F, Fondazione I, Lucia S, Neuroscienze D, Spaziale B, Vergata T. Internal Models of Target Motion[⍰]: Expected Dynamics Overrides Measured Kinematics in Timing Manual Interceptions.

